# Genomic Variability of the HCT116 Cell Line Identified Using Oxford Nanopore Sequencing

**DOI:** 10.64898/2026.04.23.720331

**Authors:** Pavel Leonov, Regina Mikheeva, Maksim Koryukov, Ekaterina Ruleva, Ekaterina Karabut, Andrey Kechin

**Affiliations:** Institute of Chemical Biology and Fundamental Medicine, Novosibirsk 630090, Russia; Novosibirsk State University, Novosibirsk 630090, Russia

**Keywords:** genomic rearrangements, structural variants, single-nucleotide variants, cell line, HCT116, colorectal cancer

## Abstract

HCT116 is a colorectal cancer cell line frequently used in anti-tumor drug development experiments as well as in studies of the molecular machinery of eukaryotic cells. It is well characterized by the presence of several single-nucleotide and short mutations in multiple oncogenes and tumor suppressor genes, including *KRAS, PIK3CA, MLH1, CTNNB1, CDKN2A, TGFBR2*, and *BRCA2*. However, its landscape of large genomic rearrangements (LGRs) and copy number variants (CNVs) is still far from being fully understood. Therefore, the aim of this study was to identify LGRs and CNVs in several HCT116 cell line samples using Oxford Nanopore sequencing technology, including three samples from the SRA NCBI database, and to compare common and unique variants across all samples. Using the recently developed eLaRodON tool, we identified 22,666 common LGRs, among which more than 70% of tandem duplications and deletions larger than 80 kb were confirmed by CNV analysis. Among LGRs affecting protein-coding sequences, two in-frame rearrangements were identified: a deletion of exons 4–6 and a duplication of exon 10 in the *CCSER1* gene, which encodes a cell division regulator protein. Given its high rearrangement rate in various tumors and the clinical significance of its overexpression, this finding may be potentially useful in future research on this cell line. Regarding differences between samples, we found that LGRs in the laboratory sample and in one of the three SRA NCBI samples occurred more frequently via ALR/Alpha repeats than via Alu repeats, in contrast to common LGRs and those unique to the other samples, a finding that may indicate the presence of unique mechanisms of genomic instability. Thus, this study reveals a broad spectrum of large genomic rearrangements and copy number variants that can be identified in the HCT116 cell line using Oxford Nanopore sequencing, including rearrangements specific to distinct cell line samples.

## Introduction

Today, more than 5,000 different tumor cell lines have been obtained from various types of cancer [1]. Their use in *in vitro* experiments with genome editing technologies or in testing new potential anti-tumor drugs allows researchers to study molecular mechanisms in normal and malignant cells, including metabolic changes and alterations in signaling pathways [2,3]. Newly developed and tested drugs can have substantially different effects on distinct cell lines, and these differences are likely associated with changes in the genetic and epigenetic context of each cell line used [3,4]. This context provides variability in gene expression and protein structure, which manifests as differences in the most important molecular processes in cells, such as cell division, proliferation, and metabolism. Therefore, new anti-tumor drugs under development must be tested on 60 cell lines, as recommended by the National Cancer Institute [5]. Consequently, a deep understanding of all single-nucleotide variants and large genomic rearrangements in the cell line used is an essential requirement for effective new drug design and for studying molecular mechanisms.

While single-nucleotide variants have been extensively studied using short-read sequencing technologies, and our understanding of their representation in cells has already reached the necessary level, the detection of large genomic rearrangements is still under active development [6]. Their identification remains challenging due to the presence of numerous extended repeat sequences in the human genome. Nevertheless, the presence of such rearrangements can substantially alter the repertoire of active genes in cells. Moreover, both tumor cells and cultured cell lines can exhibit significant heterogeneity, including with respect to large genomic rearrangements [7,8].

The HCT116 cell line was obtained from a male patient with colorectal cancer in 1981 [9] and has since been distributed to various research groups worldwide, undergoing many passages. Consequently, different samples of this cell line can substantially differ between research groups, and this variability can influence the results of *in vitro* experiments using these cells. While the NA12878 (HG001) cell line has gold-standard lists of structural variants and single-nucleotide variants [10], for the HCT116 cell line, only COSMIC copy number alterations and structural variants are known, which obviously do not cover the full spectrum of these genetic changes. At the same time, Oxford Nanopore sequencing technology, like other similar techniques (CycloneSEQ and Qitan Tech), can produce significantly longer sequencing reads and thus provides an opportunity to uncover large genomic rearrangements mediated by extended repeat sequences [11]. Recently, the authors developed a new bioinformatics tool, eLaRodON (https://github.com/aakechin/eLaRodON), which can identify more large genomic rearrangements than other existing tools and allows detection of rearrangements supported by only one sequencing read. Therefore, the aim of this study was to compare the genome structures of several HCT116 cell line samples and to identify genetic variants that are common to all samples studied, as well as those that could potentially explain some metabolic differences between these cells or their responses to chemical drugs.

## Materials and Methods

### Cell Culture and DNA Isolation

The experiments were performed using the HCT116 cell line, obtained from the collection of the Institute of Cytology and Genetics, Siberian Branch of the Russian Academy of Sciences (Novosibirsk, Russia). Mycoplasma-tested HCT116 cells were maintained in Dulbecco’s Modified Eagle’s Medium supplemented with 10% (v/v) heat-inactivated fetal bovine serum, 2 mM L-glutamine, 100 U/mL penicillin, and 100 μg/mL streptomycin under standard conditions at 37 °C in a humidified atmosphere containing 5% CO□.

Genomic DNA was extracted from cells using the SolPure HW DNA extraction kit (Magen Biotechnology Co., Ltd., Guangzhou, China) according to the manufacturer’s protocol for genomic DNA isolation from cultured cell lines. DNA quality was assessed using an ND-100C spectrophotometer (Hangzhou Miu Instrument Co., Ltd, China), and DNA concentration was quantified using a Qubit fluorometer (Thermo Fisher Scientific, Waltham, MA, USA) with the SynQuant BR DNA-100 reagent kit (Syntol, Moscow, Russia) in accordance with the manufacturer’s protocol.

### DNA Library Preparation and Sequencing

Sequencing libraries were prepared using the NEBNext Ultra II kit (New England Biolabs, Ipswich, MA, USA) and the ligation sequencing DNA kit V14 (SQK-LSK114) according to the manufacturers’ protocols. The input DNA concentration was 291 ng/µL, yielding a final amount of 1 µg per reaction. Sequencing was performed on the MinION Mk1B platform (Oxford Nanopore Technologies, Oxford, UK) using a R10.4.1 flow cell (FLO-MIN114) following the manufacturer’s protocol. Two sequencing runs were conducted, each lasting 72 hours.

### External Sequencing Data

For comparative analysis, publicly available whole-genome sequencing datasets were retrieved from the NCBI Sequence Read Archive (SRA). ONT sequencing data for the HCT116 cell line were obtained under accession numbers SRR27935354, SRR27935355, and SRR27935356 (BioProject PRJNA1075154). Raw FASTQ files were downloaded using the SRA Toolkit [12] and processed using the same bioinformatics pipeline described below to ensure methodological consistency across all comparisons. For independent validation of the identified point variants, an Illumina whole-genome sequencing dataset for the HCT116 cell line was obtained from the NCBI SRA under accession number SRR8639145 (BioProject PRJNA523380).

### Sequencing Data Analysis

After sequencing, base calling was performed using Dorado (version 1.3.0; Oxford Nanopore Technologies, Oxford, UK) with default parameters. Reads were subsequently aligned to the human reference genome version GRCh38.p14 using minimap2 (version 2.24-r1122) [13]. Point mutations were called with the Clair3 tool [14]. Large genomic rearrangements were identified with eLaRodON (https://github.com/aakechin/eLaRodON) and filtered based on the number of supporting reads (at least three supporting reads in at least one of the samples), variant allele frequency (at least 0.3 in at least one of the samples), and proximity to the hg38 genome assembly gaps. Any LGR located within 100 bp of an assembly gap was filtered out. Copy number variations were called using mosdepth [15] and ONT-spectre (https://github.com/nanoporetech/ont-spectre) tools. Sequencing data quality was assessed using samtools (version 1.23.1) [16], Nanoplot (version 1.46.2) [17], and FastQC (version 0.12.1).

The Illumina sequencing data were mapped to the human reference genome using BWA-MEM2 [18], and variant calling was also performed using Clair [14] with parameters for paired-end Illumina data. To determine the impact of the variants on the encoded protein sequence, the snpEff tool was used [19].

### Comparison of Large Genomic Rearrangements and Copy Number Variations

To identify LGRs common to all the HCT116 samples, the VCF files produced by eLaRodON were compared according to the following rules. Two variants were considered identical across samples if their types and junction directions matched, and if their genomic coordinates and sizes differed by no more than 10% or 200 bp (whichever was more stringent); for inter-chromosomal rearrangements (BND_TRL), the positional tolerance was extended to 500 bp. Such threshold was chosen based on the possible inaccuracy in the alignment of large reads.

To identify LGRs corresponding to copy number variations (CNVs), we attempted to find deletions or tandem duplications with the same or broader (but with a size difference of less than 500,000 bp compared to the CNV size) coordinates and with a positional difference relative to the CNV of less than 400,000 bp. These parameters were determined by comparing CNVs for which LGRs with a confident number of supporting reads (at least three) had been identified. The results are presented in **Supplementary Figure S1**.

For the analysis of junction boundaries, an LGR was considered to arise through repeat sequences if the two repeat sequences at the boundaries were identical.

### Statistical Analysis

For statistical analysis, the SciPy [20] and NumPy [21] Python modules were used. To test the statistical significance of differences in the ratios of the number of LGRs between samples, Fisher’s exact test with Bonferroni correction for multiple comparisons was applied.

## Results

### Sequencing yield and NGS reads statistics

Whole-genome ONT sequencing of the merged dataset from two ONT runs for the laboratory’s HCT116 cell line sample generated a total of 8,759,076 reads, yielding 24.16 Gb of sequence data with a read length N50 of 14,051 bp (mean read length: 2,758 bp; standard deviation: 7,451 bp) (**Table 1**). The mean basecall quality score was Q15.3 (median Q18.7), with 93.6% of reads exceeding Q10. The mean genome-wide coverage was 7.16×, which, while below the threshold recommended for SNV calling, is sufficient for split-read-based SV detection at the read lengths obtained. Per-autosome coverage ranged from 5.93× (chr15) to 9.61× (chr8) for the laboratory’s sample, and from 6.59× to 14.96× for the other samples, with similar per-chromosome values across samples (**Figure 1**). Coverage values for chromosomes X and Y were 3.65× and 0.57×, respectively, which is consistent with the male origin of the HCT116 cell line.

**Table 1.** Summary of sequencing metrics for the HCT116 cell line samples analyzed in this study. Lab – the laboratory sample of the HCT116 cell line. 54 – SRR27935354, 55 – SRR27935355, 56 – SRR27935356.

| **Metrics** | **Lab** | **54** | **55** | **56** | **SRR8639145** |
| --- | --- | --- | --- | --- | --- |
| Sequencing Platform | ONT | ONT | ONT | ONT | Illumina |
| Total Reads, millions | 8.76 | 3.20 | 2.00 | 2.85 | 640.98 |
| N50, bases | 14051 | 13302 | 15245 | 13767 | 101 |
| Mean Read Length | 2758 | 11268 | 11984 | 11139 | 93 |
| Mapping rate, % | 100.00 | 99.96 | 99.95 | 99.98 | 97.84 |
| Total Yield, Gb | 24.16 | 29.9 | 20.3 | 27.2 | 59.8 |
| Mean Coverage | 7.16 | 11.28 | 7.62 | 10.2 | 17.7 |
| GC Content, % | 41.0 | 41.3 | 41.4 | 41.5 | 43.2 |

**Figure 1.**
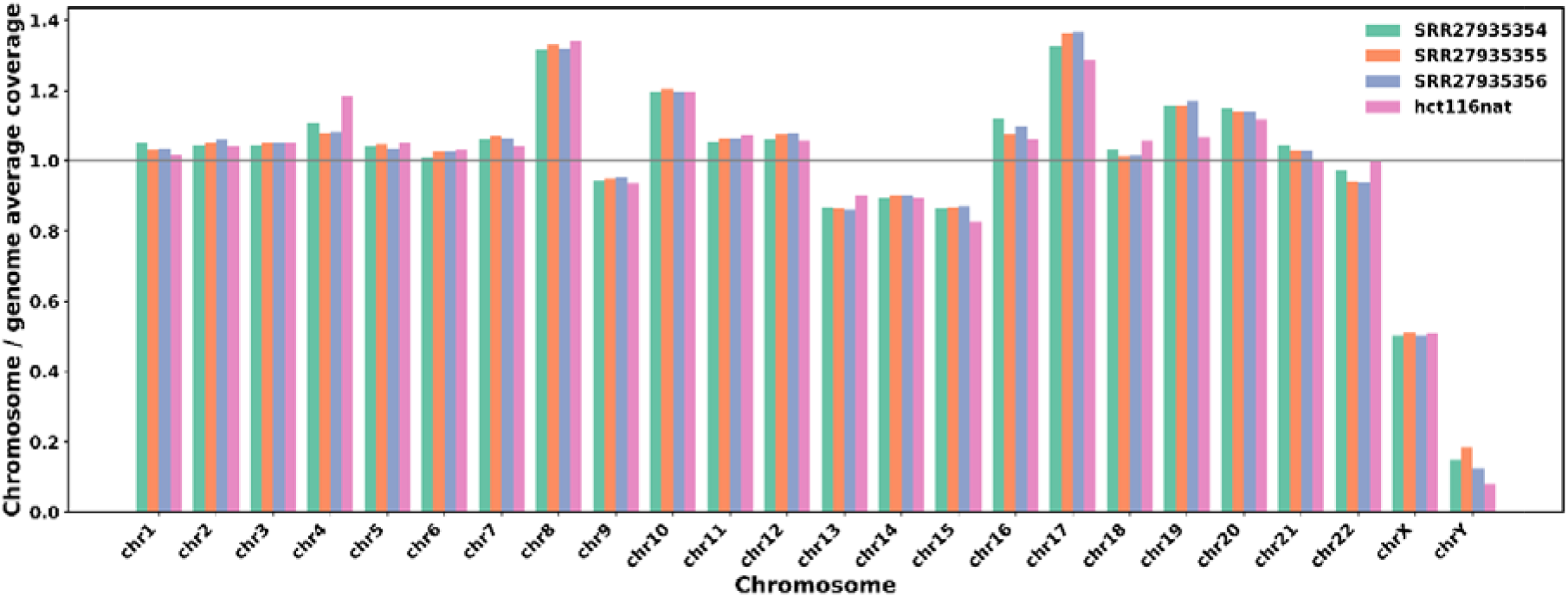
Ratio of mean chromosome coverage to mean genome coverage for all chromosomes of the samples sequenced with Oxford Nanopore technology.

### Single Nucleotide Variants and Short Insertions and Deletions

To identify SNVs and short InDels common to all samples and sample-specific ones, we applied Clair3 which yielded a total of 5459060–7022823 raw point mutations and short insertions and deletions (InDels) per sample with minimal value for Illumina sequencing data. Considering that the Illumina sequencing technology has a lowest error rate, we filtered out the Illumina’s variants with the following thresholds: number of reads with alternative allele ≥ 6, variant allele frequency (VAF) ≥ 0.2. Such filtering left 4,480,712 variants for the subsequent analysis. Among them, 3,735,607 variants were identified simultaneously in all HCT116 samples studied with 11,559 variants probably affecting proteins’ amino acid sequences with 1,557 written in the COSMIC database for the HCT116 cell line (**Supplementary Table S1**).

Among the common variants all the known point pathogenic mutations for the HCT116 cell line were successfully identified in almost all samples. The canonical oncogenic driver mutations characteristic of HCT116 – *KRAS* p.G13D (c.38G>A), *PIK3CA* p.H1047R (c.3140A>G), *CTNNB1* p.S45del (c.133_135delTCT), biallelic inactivation of *MLH1* (c.755C>A, p.S252*), frameshift alterations in *TGFBR2* (c.383delA, p.Lys153fs), *CDKN2A* (c.97delG p.Glu33fs and c.68dupG p.Arg24fs), and *BRCA2* (c.8021dupA, p.I2672fs). Only *BRCA2* c.8021dupA was not identified in the SRR27935356 sample due to the low coverage value (three reads) in that region. At the same time, mutations in the *TGFBR2* and *CDKN2A* have not been written in the COSMIC database (version 103) for the HCT116 cell line.

### Large genomic rearrangements

To investigate large genomic rearrangements in the HCT116 cell line samples, we began by examining their karyotypes and chromosomal abnormalities exceeding 10 Mb in size. The karyotype of the HCT116 cell line has been previously described with varying levels of detail. In the review by Turid Knutsen and colleagues, it was reported as 45,X,Y,der(4)t(4;17)(q3?;?q21), del(9)(q11), der(10)dup(10)(q23.1q26.1)t(10;16)(q26.1;q23), der(16)t(8;16)(q13;pter), der(18)(:4q3?::17q?22→17q21.3::18pter→18qter) [22]. In the study by Alisa Morshneva and coauthors, the HCT116 cell line was reported to have a 46,XY,der(10)dup(10)(q22q23),der(8),t(8;16)(q13;p13.3),der(18)t(17;18)(q11.2;p11.2) karyotype [23]. Oxford Nanopore sequencing technology theoretically provides an opportunity to identify the exact boundaries of genomic rearrangements. Therefore, we attempted to identify the listed rearrangements among the LGRs and CNVs detected. Among four interchromosomal translocations, only one was identified with exact boundaries as BND_TRL, supported by 3–6 reads (**Table 2**). For large chromosomal duplications and deletions, CNV boundaries were determined only using the ONT□spectre tool without confirmation with eLaRodON.

**Table 2.**
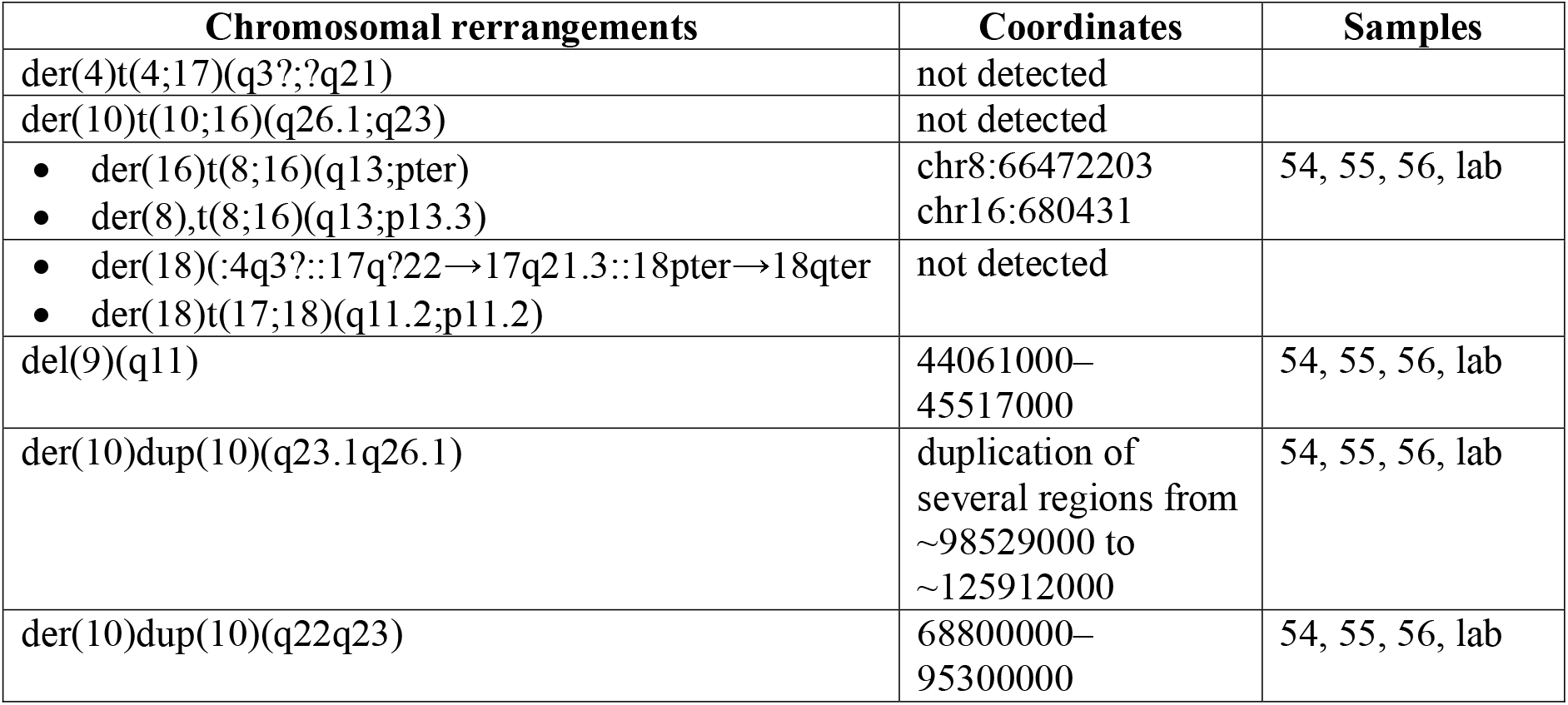
Large chromosomal rearrangements and CNVs identified with eLaRodON and/or ONT-spectre tools.

In addition to large chromosomal rearrangements, smaller deletions and duplications ranging in size from tens of thousands to several million base pairs were identified using ONT-spectre, some of which were confirmed by LGR boundaries from eLaRodON (**Table 3**). Although ONT-spectre yielded more CNVs than LGRs of similar size, including both common and sample-specific CNVs (**Supplementary Table S2**), confirmation of these events will require higher depth of human genome coverage or the application of alternative methods such as droplet digital PCR. For chromosomes with highest mean coverage (chr17, chr8, and chr10), the number of duplicated genomic regions was also the highest (**Figure 1** and **Supplementary Table S2**).

**Table 3.** Copy number variations (CNVs) identified in all HCT116 cell line samples, some of which were confirmed as large genomic rearrangements (LGRs) using the eLaRodON tool. CNVs previously reported in the COSMIC database are highlighted in bold. Coordinates of CNVs specified from the COSMIC database are shown in italic.

| Chr | Start | End | Length | Type | Zyg. | Samples | Method |
| --- | --- | --- | --- | --- | --- | --- | --- |
| chr2 | 140940220 | 141441937 | 501,718 | Del | Hetero | All | LGR, CNV |
| chr2 | 204625353 | 204930317 | 304,965 | Del | Hetero | All | LGR, CNV |
| chr3 | 60235172 | 60404776 | 169,605 | Del | Hetero | All | LGR, CNV |
| chr3 | 60604285 | 60843856 | 239,572 | Del | Hetero | All | LGR, CNV |
| chr3 | 91555000 | 91713000 | 158,000 | Del | Homo | All | CNV |
| chr4 | 90391268 | 90690995 | 299,728 | Del | Hetero | All | LGR, CNV |
| chr6 | 1783057 | 2034965 | 251,909 | Del | Hetero | All | LGR, CNV |
| chr6 | 2030628 | 2268487 | 237,860 | Del | Hetero | All | LGR, CNV |
| chr6 | 20375926 | 20624595 | 248,670 | Dup | Hetero | All | LGR, CNV |
| chr6 | 58447373 | 58798642 | 351,270 | Del | Homo | All | LGR, CNV |
| chr16 | 6165928 | 7090949 | 925,022 | Del | Hetero | All | LGR, CNV |
| **chr16** | **6172438** | **6884622** | **712,185** | **Del** | **Homo** | **All** | **LGR, CNV** |
| chr16 | 78228812 | 78442742 | 213,931 | Del | Hetero | All | LGR, CNV |
| **chr17** | **18455633** | **18561273** | **105,641** | **Del** | **Homo** | **All** | **CNV** |
| chr17 | 24250000 | 24899000 | 649,000 | Del | Homo | All | CNV |
| chr17 | 25015000 | 26509000 | 1,494,000 | Del | Homo | All | CNV |
| chr19 | 20490322 | 20841029 | 350,708 | Del | Hetero | All | LGR, CNV |
| chr19 | 53990998 | 55113922 | 1,122,925 | Del | Hetero | All | LGR, CNV |
| chr20 | 29865027 | 30945361 | 1,080,335 | Del | Homo | All | LGR, CNV |
| chr22 | 18204639 | 18893221 | 688,583 | Del | Homo | All | LGR, CNV |
| chrX | 29364601 | 29501641 | 137,041 | Dup | Homo | All | LGR, CNV |
| **chrX** | **29523494** | **29945784** | **422,294** | **Del** | **Homo** | **All** | **LGR, CNV** |

The next group of LGRs comprised variants identified with eLaRodON simultaneously in all samples as well as sample□specific variants (**Supplementary Tables S3–S7**). The total number of common LGRs of size 50 bp or larger was 22,666 variants (8,723 deletions, 248 tandem duplications, 52 inversions, 138 interchromosomal translocations, and 13,505 insertions). Some of these were confirmed by CNV analysis using ONT□Jspectre or by depth□of□coverage analysis using mosdepth. Three BND_DEL rearrangements larger than 2–27 Mb were not confirmed by coverage analysis, suggesting possible translocation of the regions instead of deletion. Although the VAF values for common LGRs were correlated between samples — in particular for LGRs with a depth of coverage of at least ten (**Supplementary Figure S3**) — high variability in the number of supporting reads and in VAF values was observed across samples. Among the LGRs common to all samples studied, 8,791 (38.8%) and 353 (1.6%) overlapped with genes and exons, respectively. The percentage of such LGRs among sample□specific variants was similar: 34–35% for SRR27935354 and the laboratory’s sample, and 44–50% for SRR27935355 and SRR27935356. Examples of three LGRs which possibly affect the structure or regulation of genes are shown in **Figure 2**. One large inversion of 18,128 bp changed the putative regulatory region of the lactate dehydrogenase D (*LDHD*) gene (about 90 Kb downstream of the gene start, **Figure 2A**). Another included a heterozygous deletion of exons 4– 6 of the *CCSER1* gene and a simultaneous tandem duplication of exon 10 with adjacent intronic regions (**Figure 2B**). Both variants did not affect the protein coding frame; therefore, a functional protein with altered properties is likely to be expressed. Considering that this gene encodes the FAM190A protein, which regulates cell division [24,25], these variants could substantially affect cellular mechanisms of mitosis.

**Figure 2.**
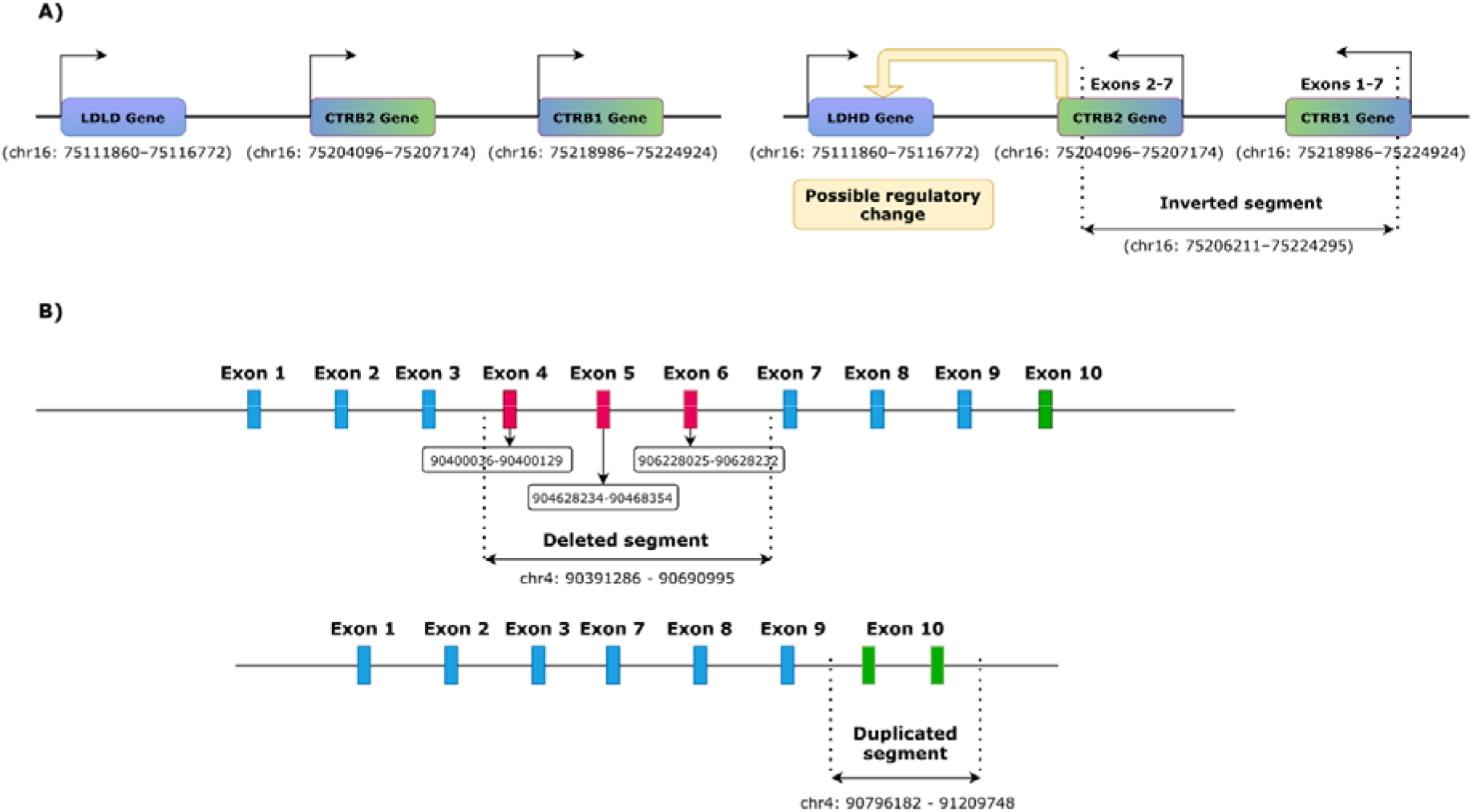
Examples of LGRs affecting possible regulatory region of lactate dehydrogenase D (*LDHD*) gene (**A**) and *CCSER1* gene encoding cell division regulatory protein (**B**).

Although all the studied HCT116 cell line samples shared many common LGRs, sample-specific variants were also identified. For example, 482 LGRs (280 insertions, 122 tandem duplications, 73 deletions, 4 inversions, and 3 interchromosomal translocations) were identified in the laboratory’s sample (**Supplementary Table S7**). Among these, 167 LGRs overlapped with genes, and 14 were likely to alter protein-coding sequences.

To assess possible differences in the mechanisms of mutagenesis, we compared the structures of common and sample-specific LGRs by their type ratios and junction boundary features, particularly the presence of microhomology or homology and the distance from repeat sequences to the breakpoint (**Figure 3**). Based on the distribution of LGR types, the most striking difference was observed between common LGRs and sample-specific ones (**Figure 3B**). Whereas LGRs present in all samples were predominantly insertions (59%) and deletions (38%), tandem duplications and interchromosomal translocations constituted a much larger share among sample-specific LGRs: 23–53% and 8–14%, respectively (p-values ranged from 2×10 □ ^23^□ to 5×10□^11^ □, Fisher’s exact test). The proportion of tandem duplications among all LGR types was also significantly lower in the laboratory’s sample compared with the other samples (p-values ranged from 2×10□^1^ □ to 5×10□^1^ □, Fisher’s exact test). Although differences in the presence of homologous sequences between junction boundaries were also observed between LGR groups, they were not statistically significant.

**Figure 3.**
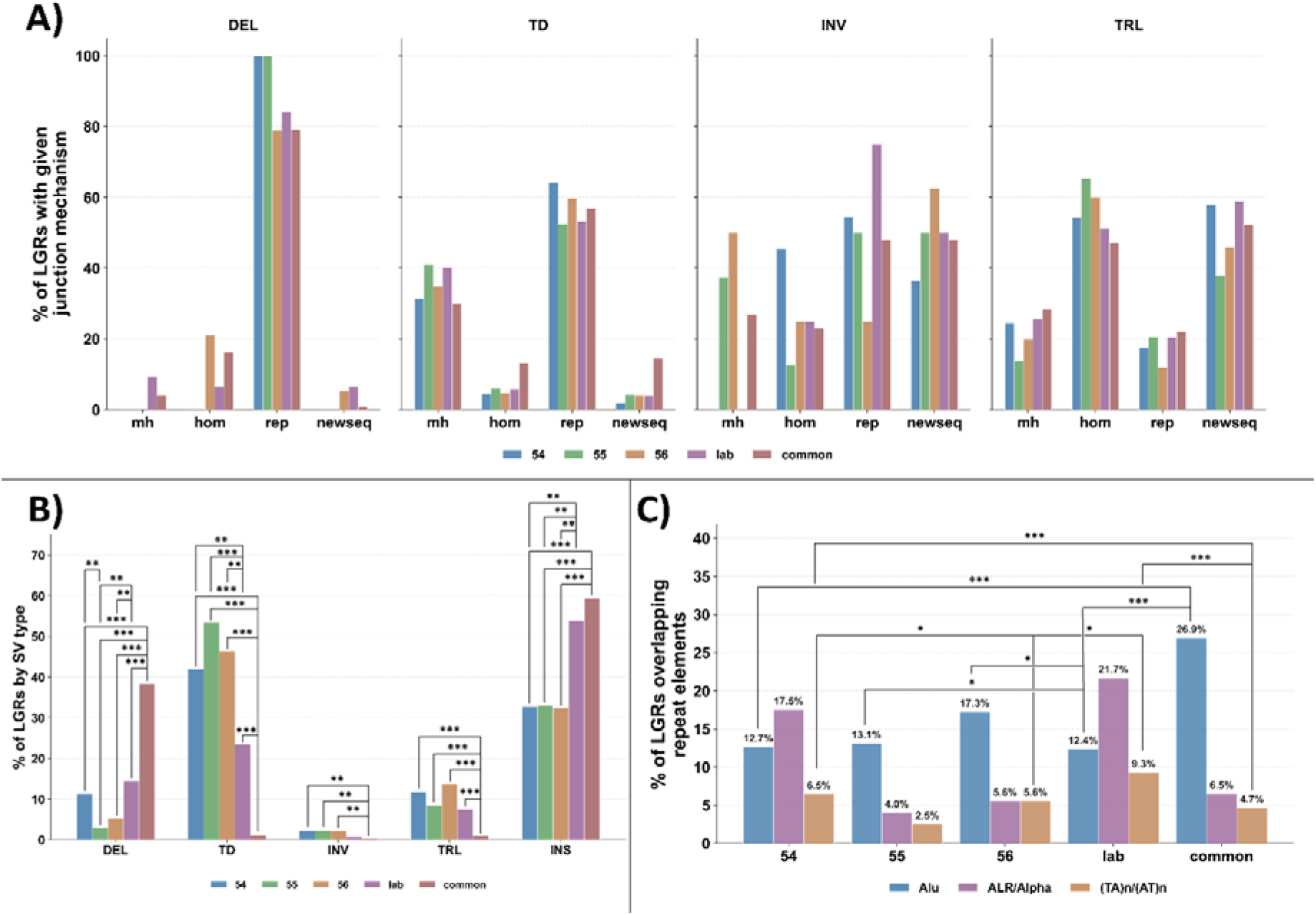
Comparison of LGRs by their ratio and type of junction boundaries. **A)** Type of junction boundary homology: microhomology (mh), homeology (hom), or repeat (rep). “Newseq” indicates the presence of a sequence of unknown origin between two joined sequences. This category does not overlap with other junction types; i.e., a novel sequence can be present together with microhomology or homeology, or in LGRs without any homology. **B)** Distribution of different types of LGRs among all samples studied. DEL – deletions, TD – tandem duplications, INV – inversions, TRL – interchromosomal translocations, INS – insertions. **C)** Distribution of LGRs by the type of repeat sequence through which the LGR occurred. Only Alu repeats, ALR/Alpha, and (TA)n with (AT)n are shown. Asterisks indicate statistical significance: * p < 0.05, ** p < 0.01, *** p < 0.001 (Fisher’s exact test with Bonferroni correction for multiple comparisons).

By LGR type ratios, the sample most similar to the laboratory’s sample was SRR27935354. This similarity was also maintained when comparing repeat sequences located at LGR boundaries (**Figure 3C**). The difference between LGRs common to all samples and sample-specific LGRs was statistically significant only for SRR27935354 and the laboratory’s sample with respect to the proportion of LGRs occurring via Alu repeats (p = 6×10□^1^ □ and 2×10□^1^ □, respectively). Thus, the similarity between SRR27935354 and the laboratory’s sample was confirmed in two independent comparisons of LGR structure.

## Discussion

In this study, we performed an in□depth comparison of several HCT116 cell line samples sequenced using Oxford Nanopore technology. By combining the results of SV and CNV detection across all samples and applying eLaRodON — a tool recently developed by the authors — we were able to identify numerous LGRs common to all studied samples, as well as several sample□specific variants. Most of these LGRs and CNVs are described for this cell line for the first time, which was made possible by the unique algorithm of eLaRodON that reports all LGRs supported by even a single sequencing read. The identification of LGRs is significantly complicated by the low probability that a sequencing read will span an LGR in a way that reaches the end of the repeat element [26,27]. If we had not included LGRs supported by only one read, we would have missed 7,710 (34%) of the LGRs common to all samples (**Supplementary Table S3**). Consequently, the number of identified LGRs would have been close to that typically reported for human DNA samples [28–32]. Moreover, many potential common LGRs were excluded from both the common and sample□specific lists because they were detected in only two or three samples. Therefore, the true number of common LGRs may be even higher than we report here.

Taking into account LGRs with only one supporting read, it would be of great interest to study subclonal LGRs, whose detection is limited not only by the complications mentioned above but also by their rare occurrence among all sequencing reads obtained [33]. Consequently, if a supporting read threshold greater than 1–2 is applied, coverage of approximately 1000× is required to detect an LGR present in only 5% of cells. Achieving such coverage would require about 200 Oxford Nanopore flow cells per sample. Nevertheless, alternative approaches are currently required to validate LGRs with single supporting reads, and the identification of subclonal LGRs will be the focus of our future research.

Although it may seem counterintuitive that we applied a higher alternative allele depth threshold for SNVs and short InDels (five or six reads) than for LGRs (three reads), this choice is explained by the higher probability of obtaining a random false□positive SNV compared to a false□positive LGR. During sequencing, the probability of an incorrectly called base occurring at the same position in two or three reads across the whole genome is much higher than that of an incorrect junction of two DNA fragments at the same chromosomal position. Therefore, we believe that our choice of thresholds is appropriate, and similar thresholds have been used in other studies [34,35].

The orthogonal confirmation of identified LGRs is a complicated task, accompanied by difficult primer design to avoid repeat sequences, and clearly cannot be carried out using only short-read sequencing techniques, as has been done in some other studies [36,37]. While large deletions and duplications can be confirmed by evaluating coverage values for the corresponding genomic regions — as was done in this study — other types of LGRs, such as inversions or translocations, require the use of other techniques. Moreover, although more than 70% of LGRs of size 80□kb or larger were confirmed by changes in depth of coverage, there were large deletions and duplications not confirmed by coverage value changes, as well as many CNVs not confirmed by LGRs. The former inconsistencies may be associated with the subclonal nature of those LGRs [38], whereas the latter may be caused by complications in LGR detection discussed above.

Among the LGRs common to all HCT116 cell line samples studied, we describe in more detail two LGRs that, in our opinion, merit attention. The first is an inversion near the lactate dehydrogenase D gene. The encoded enzyme catalyzes the conversion of D-lactate to pyruvate, as well as other D-2-hydroxyacids [39]. Its elevated level in colorectal cancer is associated with poor overall survival [40], and changes in its expression can be caused by various molecular mechanisms, including the potential ones identified here for the HCT116 cell line. The second LGR is a complex rearrangement: an in-frame large deletion of three exons accompanied by an in-frame duplication of another exon in the *CCSER1* gene, which encodes a cell division regulator [24,25]. Deletions of certain exons of this gene have already been described in the literature and may lead to its hyperexpression, thereby enhancing proliferation and genomic instability [25]. Other studies, however, suggest that altered CCSER1 could be considered a target neoantigen for immunotherapy [41]. Nevertheless, to the best of our knowledge, a dual rearrangement of the type identified in our study has not been reported previously.

A potentially useful difference between the cell line samples was identified regarding the junction boundary structure of sample-specific LGRs. For two samples — the laboratory sample of the HCT116 cell line and SRR27935354 — we observed a prevalence of LGRs mediated by ALR/Alpha repeat sequences rather than Alu repeats, in contrast to what was observed for the other samples and for common LGRs. This suggests that additional mechanisms of genomic instability, beyond the known mismatch-repair deficiency, may have been acquired in these two samples during previous passages. Given that ALR/Alpha repeats are predominantly located in centromeric and pericentromeric regions, a higher rate of aneuploidy and chromosomal instability could be expected for these samples [42,43].

