## Supplementary Figure for "Genomic Variability of the HCT116 Cell Line Identified Using Oxford Nanopore Sequencing"

**Supplementary Figure S1.** Comparison of differences in position and structural variant length for copy number variations (CNVs) for which corresponding large genomic rearrangements (LGRs) with at least three supporting reads were identified (green dots), and CNVs for which LGRs with only one or two supporting reads were found (red dots). The horizontal axis represents the difference in the length of the mutated region between a CNV and an LGR. The vertical axis represents the maximal difference between the start or end positions of a CNV and an LGR.

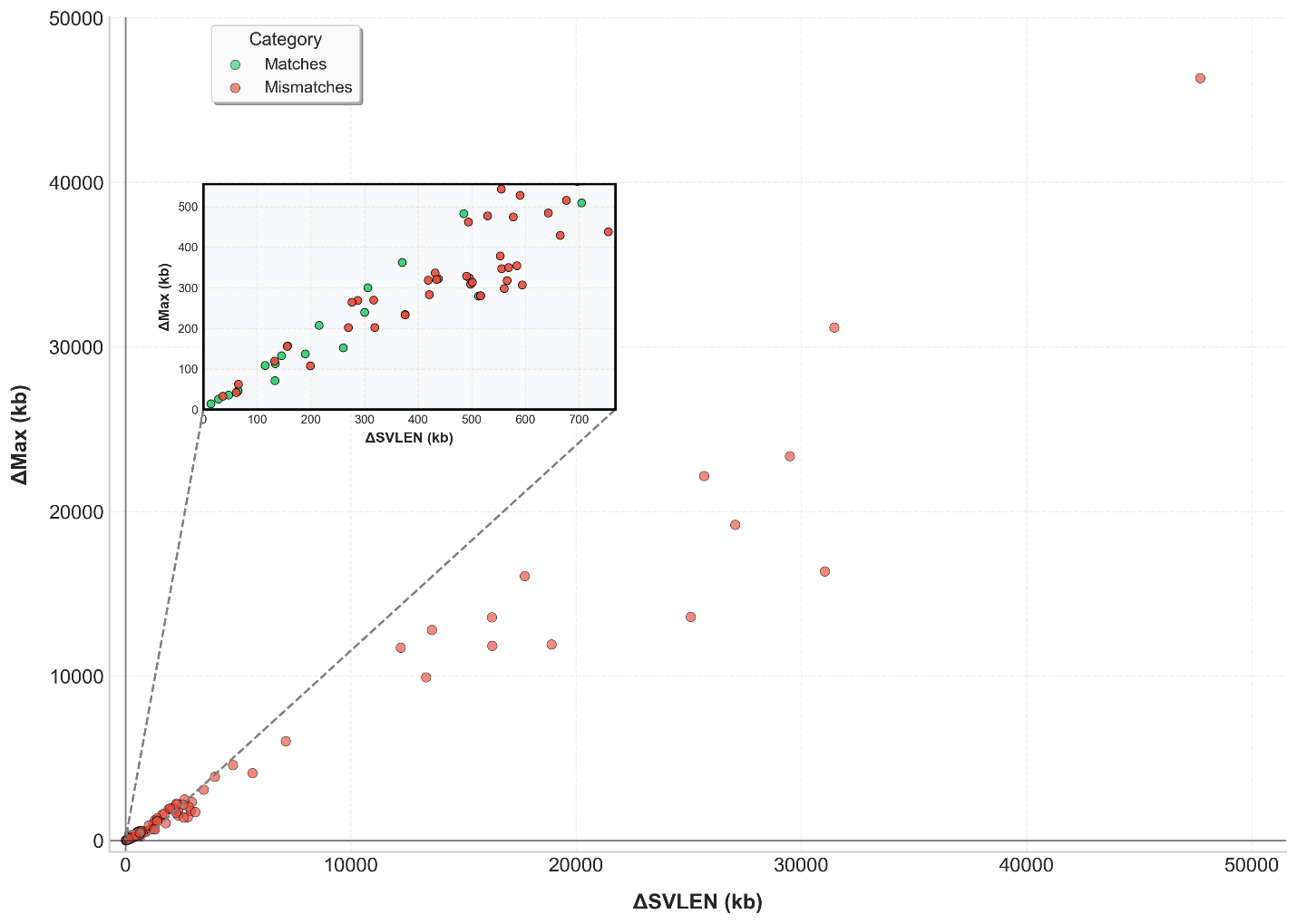

**Supplementary Figure S2.** Pair-wise comparisons of variant allele frequency values for large genomic rearrangements (LGRs) common to all HCT116 cell line samples studied. Only LGRs with DP ≥ 10 in both samples are shown.

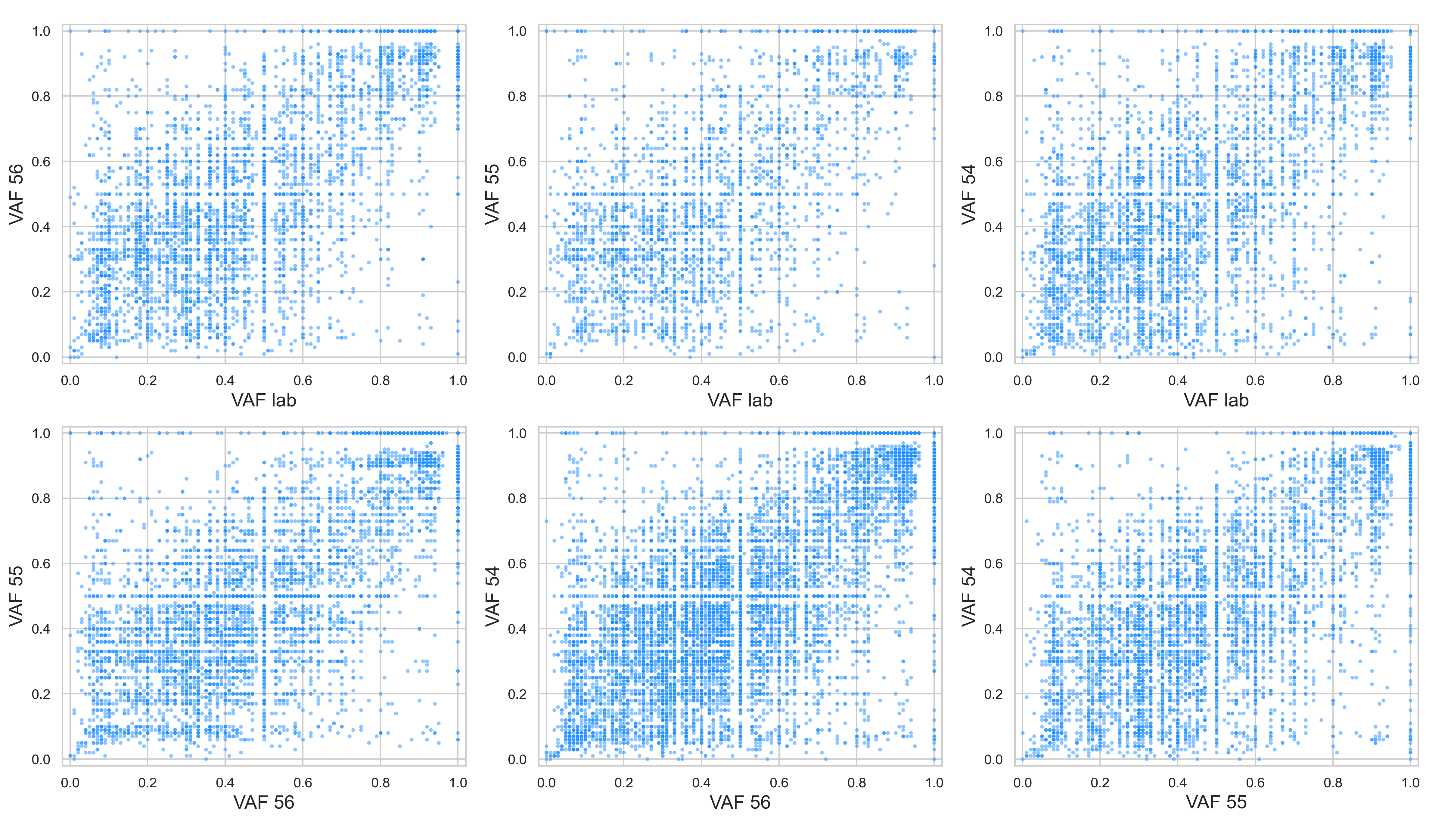
